# Feasibility of automated insulin delivery guided by continuous glucose monitoring, in extremely preterm infants

**DOI:** 10.1101/374801

**Authors:** Kathryn Beardsall, Lynn Thomson, Daniela Elleri, David B Dunger, Roman Hovorka

## Abstract

**One Sentence Summary:** Closed loop systems based on subcutaneous glucose measurements could provide an efficacious and safe means of optimizing glucose control in preterm infants while reducing resources required including time of bedside personnel.

**Abstract:** Closed loop systems have been used to optimise insulin delivery in children with diabetes, but they have not been tested in neonatal intensive care. Extremely preterm infants are prone to life-threating hyperglycaemia and hypoglycaemia; both of which have been associated with adverse outcomes. Insulin delivery is notoriously variable in these babies and time-consuming, with management requiring frequent changes of dextrose-containing fluids and careful monitoring. We aimed to evaluate the feasibility of closed loop management of glucose control in preterm infants in a single centre feasibility study with a randomised parallel design. Eligibility criteria included birth weight <1200g and <48hours of age. All infants had subcutaneous continuous glucose monitoring for the first week of life, with those in the intervention group receiving closed loop insulin delivery between 48 and 72hours of age. The primary outcome was percentage of time in target (sensor glucose 4-8mmol/l). The mean (SD) gestational age and birth weight of intervention and control study arms were 27.0(2.4) weeks, 962(164) g and 27.5(2.8) weeks, 823(282) g respectively. The time in target was dramatically increased from median (IQR) 26%(6,64) with paper guidance to 91%(78, 99) during closed loop (p<0.001), without increasing hypoglycaemia. There were no serious adverse events and no difference in total insulin infused. Closed loop glucose control based on subcutaneous glucose measurements is feasible and appears to provide an efficacious means of optimising glucose control in extremely preterm infants.

## Introduction

Preterm infants are at high risk of both hyperglycaemia and hypoglycaemia, predominantly related to deficit of insulin production and glycogen stores, with additive effects of parenteral nutrition, inotropic drug infusions and sepsis related insulin resistance.(*1, 2*) Hyperglycaemia is observed in 80% of preterm infants, and glucose variability is associated with increased mortality and morbidity.(*2-6*) Moreover, at a practical level glucose control is difficult to achieve in an extremely low birth weight preterm infant; often this requires multiple changes on intravenous infusions and insulin dosing requiring extra attention of bedside staff and general expense. The use of sliding scale insulin therapy is widespread but considered suboptimal as the desire to minimize blood sampling, (*7*) combined with the extremely variable response to insulin,(*1*) puts these babies at risk from hypoglycaemia.(*8*) This results in babies often being managed with a reduction in parenteral nutrition, and potentially inadequate nutritional support, at a critical time of growth and development.

Continuous glucose monitoring has been used in neonatal intensive care to identify hypoglycaemia and is considered sufficiently accurate to support clinical management.(*8, 9*) (*10*) However, the wide variation in individual insulin sensitivity and the limited staff resources make it challenging for the full potential of continuous glucose monitoring to be realized. Adaptive computerized algorithms utilizing hourly to four hourly blood glucose measurements have been evaluated in adults (*11-13*) and neonates undergoing intensive care. (*10*) (*14*) The addition of frequent glucose levels obtained by continuous glucose monitoring allows the development of closed loop systems as investigated in adult intensive care patients documenting its safety and efficacy.(*15, 16*) The present study hypothesized that closed loop based on subcutaneous continuous glucose monitoring can be similarly effective in informing insulin delivery and targeting glucose control in extremely preterm infants compared with continuous glucose monitoring with insulin therapy guided by a paper algorithm.

## Results

### Study Population

Ten babies were randomly assigned to closed loop intervention and ten babies were randomly assigned to continuous glucose monitoring with insulin therapy guided by a paper algorithm. Baseline characteristics of the two groups were similar (Table 1).

**Table 1.** Baseline demographic data.

|  | Closed loop | Control |
| --- | --- | --- |
|  | (n=10) | (n=10) |
| Gestational age at birth (week) | 27.0 (2.4) | 27.5 (2.8) |
| Birth weight (g) | 962 (164) | 823 (282) |
| Sex (male:female) | 5:5 | 5:5 |
| **Antenatal variables** |  |  |
| Antenatal steroids | 10 (100%) | 9 (90%) |
| Maternal smoking | 1 (10%) | 2 (20%) |
| Chorioamnionitis | 2 (20%) | 3 (30%) |
| PROM<sup>a</sup> | 4 (40%) | 4 (40%) |
| Hypertension | 1 (10%) | 1 (10%) |

<sup>a</sup>PROM: prolonged rupture of membranes (>24hours)
Data are presented as mean (SD)

All 20 babies remained in the study throughout the intervention period from 48 to 72h with comparable amount of sensor glucose data available for analyses in each group [median (IQR) for both study groups 24h (23.75, 24.00)]. Control algorithm directed insulin therapy was followed at all times during the pre-specified 48-72h period. The maximum period of sensor signal loss during the closed loop was 3.5h during which hourly blood glucose values were used by the control algorithm.

### Glucose control, insulin and dextrose administration between 48 and 72 hours

There was no difference in the baseline mean sensor glucose at 48h between study groups (Table 2). During the period 48 to 72 hours, the time spent in the target glucose range between 4.0 and 8.0mmol/l, the primary endpoint, was significantly higher in babies in the closed loop group (91% (78, 99); median (IQR)) compared to controls [26% (6, 64); p<0.001]. Similarly, the time spent in the wider target range 2.6 to 10.0mmo/l was higher in the closed loop group with median 100% compared to control group with median 84% (p=0.03). This was predominantly related to the smaller percent of time with sensor glucose values above10mmol/l with median 16% in the control group compared to median 0% in the closed loop group. There was no difference in the time spent with sensor glucose levels less than 2.6mmol/l. Lower sensor glucose was observed in the closed loop group median (IQR) 6.2 (6.1, 7.1)mmol/l compared to the control group 8.6 (7.4, 11.1)mmol/l (p=0.002). Glucose variability as measured by the standard deviation of sensor glucose was similar (p=0.604).

**Table 2.** Comparison of glucose control, insulin delivery, and nutritional intake during the intervention period (48 to 72 hours post birth).

|  | **Closed-loop** | **Control** | **p-value** |
| --- | --- | --- | --- |
|  | **(n=10)** | **(n=10)** |  |
| Time spent with sensor glucose level (%) |  |  |  |
| 4.0 to 8.0 mmol/l<sup>a</sup> | 91 (78, 99) | 26 (6, 64) | <0.001 |
| 2.6-10 mmol/l | 100 (94,100) | 84 (46, 98) | 0.133 |
| > 10.0 mmol/l | 0 (0, 6) | 16 (2, 54) | 0.113 |
| < 2.6 mmol/l | 0.0 (0.0, 0.0) | 0.0 (0.0, 0.0) | 0.720 |
| Baseline sensor glucose (mmol/l) | 7.9 (6.9, 11.5) | 8.2 (7.0, 12.4) | 0.182 |
| Mean sensor glucose (mmol/l) | 6.2 (6.1, 7.1) | 8.6 (7.4, 11.1) | 0.002 |
| SD of sensor glucose (mmol/l) | 1.0 (0.8, 1.9) | 1.3 (0.9, 2.5) | 0.604 |
| Episodes of blood glucose <2.6 mmol/l<sup>b</sup> | 1 | 0 | 1.000 |
| Insulin (U/kg/hour) | 0.04 (0.03, 0.07) | 0.02 (0.00, 0.11) | 0.400 |
| Nutritional intake |  |  |  |
| Dextrose (mg/kg/min) | 8.4 (7.2, 10.3) | 8.5 (4.2, 10.6) | 0.604 |
| Protein (g/kg/day) | 3.2 (2.5, 4.1) | 3.5 (1.6, 4.1) | 1.000 |
| Lipid (g/kg/day) | 1.8 (1.0, 1.8) | 1.4 (0.9, 2.2) | 0.905 |
| Trophic feeds | 4 | 4 | 1.000 |

<sup>a</sup>Primary endpoint
<sup>b</sup>Present at start of closed loop study period prior to computer algorithm advice being initiated
Data are presented as median (interquartile range)

The summative glucose profiles for each study group as well as insulin infused are provided in Figure 1. Nine out of the ten babies in the closed loop group had received insulin prior to the 24 hour intervention with the one remaining baby being started on insulin during the 24 hour closed loop. This compared to four babies in the control arm having received insulin prior to, and eight babies receiving insulin during the intervention period. Four babies in the closed loop study arm received additional 20% dextrose for short periods during the intervention period (up to 3.5 hours). The mean infusion rate in these babies ranged from 0.13ml/kg/hour to 0.53ml/kg/hour. The highest rate being infused in a baby who was hypoglycaemia prior to the start of the closed loop intervention. There was no statistical difference in the total amount of insulin infused or nutritional intake between study groups during the 24 hour intervention period.

**Figure 1.**
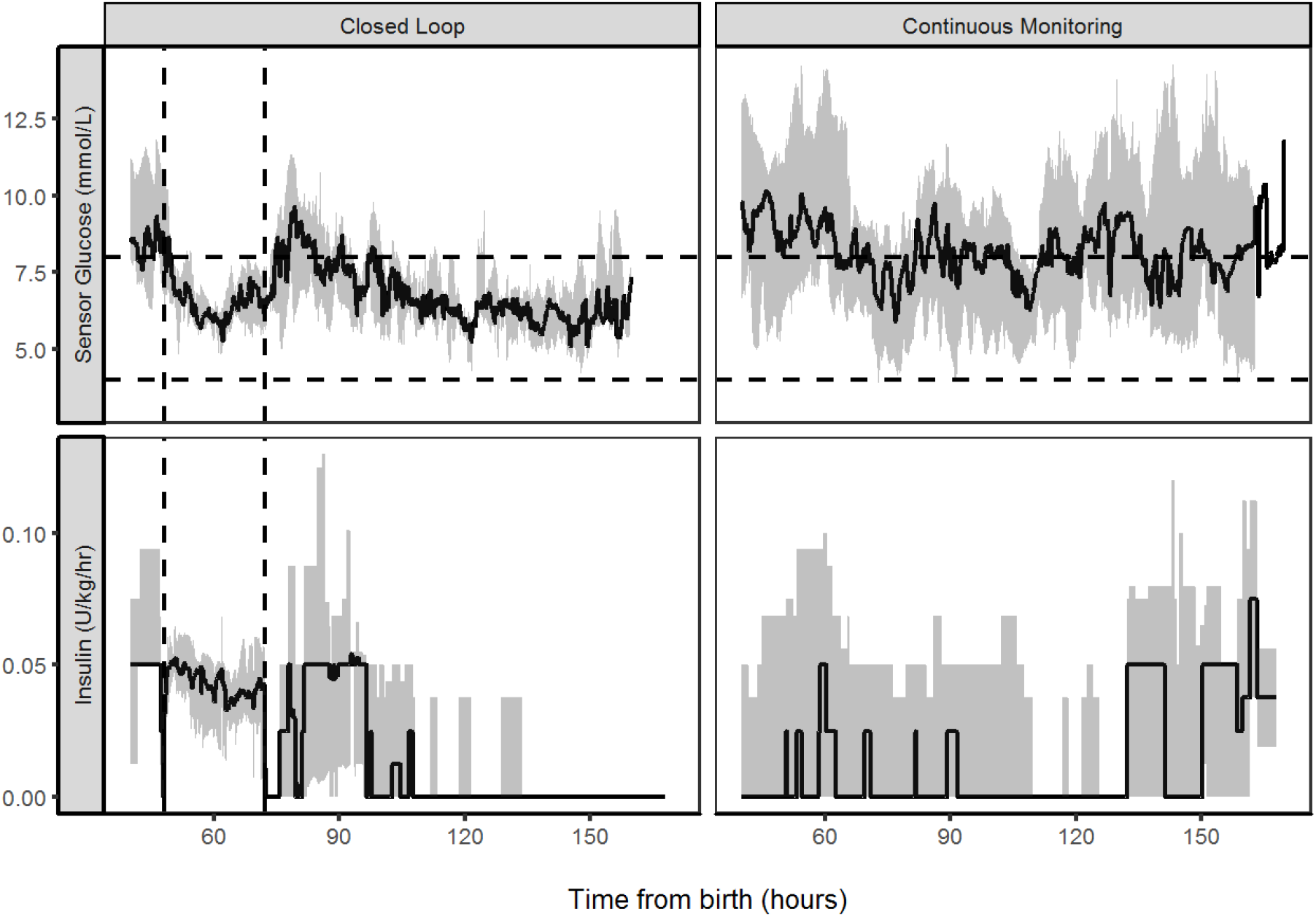
Glucose control and insulin delivery. Median (IQR) of sensor glucose and insulin infused in babies randomized to closed loop management or continuous glucose monitoring with paper algorithm (control). The closed loop intervention period is denoted by the vertical lines, and the target glucose range 4.0-8.0mmol/l is denoted by horizontal lines.

### Glucose control, insulin and dextrose administration between 72 and 160 hours

In the post intervention period after 72 hours, there was a trend of increased time in both glucose target ranges (4.0-8.0mmo/l and 2.6-10.0mmol/l) in the closed loop group compared to the control group (Table 3), but these differences did not reach statistical significance.

**Table 3.** Comparison of glucose control, insulin delivery, and nutritional intake during the post intervention period (72 to 160 hours post birth).

|  | Closed-loop | Control | p-value |
| --- | --- | --- | --- |
|  | (n=10) | (n=10) |  |
| Time spent with sensor glucose level |  |  |  |
| 4.0 to 8.0mmol/l | 64 (39, 90) | 42 (29.67, 55) | 0.053 |
| 2.6-10mmol/l | 95 (79, 97) | 78 (61.45, 97) | 0.243 |
| >10mmol/l | 3 (1, 21) | 22 (3, 36) | 0.156 |
| <2.6mmol/l | 0.0 (0.0, 0.8) | 0.0 (0.0, 0.2) | 0.720 |
| Mean sensor glucose (mmol/l) | 7.0 (6.8, 8.5) | 8.3 (7.3, 9.2) | 0.182 |
| SD of sensor glucose (mmol/l) | 1.7 (1.5, 2.1) | 1.7 (1.3, 2.8) | 0.780 |
| Episodes of blood glucose <2.6mmol/l | 1 | 0 | 1.000 |
| Insulin (U/kg/h) | 0.02 (0.00, 0.04) | 0.03 (0.00, 0.05) | 0.356 |
| **Nutritional intake** |  |  |  |
| Dextrose (mg/kg/min) | 9.4 (7.0, 10.6) | 8.7 (5.4, 10.9) | 0.549 |
| Protein (g/kg/day) | 3.7 (2.7, 4.3) | 3.8 (2.9, 4.4) | 0.968 |
| Lipid (g/kg/day) | 2.0 (1.5, 2.9) | 1.8 (1.2, 2.8) | 0.611 |
| Oral milk intake (ml/kg/day) | 4.4 (3.5, 11.5) | 5.0 (0.5, 13.0) | 0.720 |

Data are presented as median (interquartile range)

### Nutrition and clinical care

All babies received parenteral and enteral nutrition according to the standard local neonatal unit protocol. During the closed loop intervention period between 48 and 72 hours four babies in each study group were receiving minimal amounts of trophic feeds, and in the post intervention period after 72 hours there was no difference in the volumes of milk received between the two study groups (Table 3).

### Safety

There were no reported concerns about the sensor site in terms of inflammation or infection either during the study or after removal. In the closed loop study group there were two babies who had documented episodes of hypoglycaemia. One episode occurred when checking the baseline blood glucose prior to the onset of closed loop, when maintenance fluids were being changed, and when no insulin was being infused. The algorithm advised 20% dextrose that was infused. The second episode was on day 6, again associated with a change of maintenance fluids. There were a further two babies who had periods (after the 72 hour closed loop) when sensor glucose fell to below 2.6mmol/l but the blood glucose at this time was documented above 2.6mmol/l, one baby had two episodes each lasting 10 minutes and one baby had a single episode lasting 25 minutes. In the control study group no babies had a documented blood glucose value less than 2.6 mmol/l. One baby in the control group had an episode lasting 205 minutes when the sensor glucose fell to less than 2.6 mmol/l (blood glucose was not checked at this time despite this being a study recommendation). None of the babies were on insulin, and there were no clinical concerns about hypoglycaemia in these babies during these episodes.

### Discussion

This study is the first to show that a closed loop system utilizing subcutaneous continuous glucose monitoring to guide insulin delivery is safe and feasible, and may improve glucose control in extremely preterm infants. This new approach could represent a step-change in care providing greater safety and tight control while minimizing staff time at bedside and changes in fluid/insulin treatment. Compared to real time continuous glucose monitoring alone, closed loop increased time when sensor glucose was in target range between 4 and 8mmol/l three fold. In the high intensity and high cost setting of neonatal intensive care, this preliminary data support further development of closed loop systems, with real-time glucose responsive insulin and dextrose delivery to support the care of these babies.

Reflecting the current controversy regarding optimal targets for glucose control in neonatal intensive care we adopted a moderate glucose target range between 4.0 and 8.0mmol/l rather than the tight glycaemic regimen of Leuven and NICE-SUGAR studies.(*17, 18*) These moderate target ranges represent physiological in utero levels(*19*), and the upper threshold reflects the postnatal glucose level which has been associated with increased mortality and morbidity in preterm infants.(*20*) The paper-based guideline for insulin therapy utilized an identical target glucose range between 4.0 and 8.0mmol/l. However, it should be noted our study does not resolve this long-standing debate in the field, but rather shows how an automated system can be used to achieve tight control to a given target range.

Different strategies are currently utilized to target glucose control in the preterm infant each with different risks and benefits. A reduction in dextrose load risks compromised nutritional intake whilst insulin therapy can lead to hypoglycaemia. In this study, the use of the continuous glucose monitoring (CGM) highlighted clinically silent episodes of hypoglycaemia in both study arms, independent of insulin use. The CGM were calibrated using the Statstrip® meter (Nova Biomedical Waltham, MA, USA) that is validated for use in the neonates and has FDA approval for use in intensive care. These meters were used, as they were the standard of care for clinical management of glucose control within the neonatal unit. Although there remains controversy, regarding the clinical significance of clinically silent episodes of hypoglycaemia detected on CGM there is recent evidence of an association between these episodes with substantially increased risk of impaired executive function and visual motor difficulty at 4.5 years.(*21*)

The frequency of blood glucose sampling in preterm infants is typically much lower than in adults and children in intensive care and previous studies have explored the potential for the use of subcutaneous continuous glucose monitoring(*10*). These studies though remained dependent on staff responding to trends or alarms in sensor glucose before intervening.(*10*). This contrasts with the present study in which the targeting of glucose control is proactively driven by the closed loop algorithm, which was responding to frequently sampled sensor glucose data. This study is unique in exploring a control approach belonging to the family of model predictive control algorithms and optimized on a validated computer simulations environment (*22*) prior to study commencement to ensure a favorable outcome.

This is the first randomized study to evaluate the feasibility of a closed loop control based on continuous glucose monitoring in preterm infants to guide insulin delivery to support glucose control. The strengths of our study are the randomized controlled study design and comparability of the study groups and nutritional intakes. The study limitations include a small sample size, and a short study duration as well as a single center study design, which limit the generalizability but do not affect the main study outcome. Further studies are required to explore the impact of a fully automated system with infusion pumps providing insulin and 20% dextrose under fully automated computer control. Thus, closed loop insulin delivery based on subcutaneous continuous glucose monitoring appears a potentially safe, feasible and efficacious approach for targeting glucose control in preterm infants requiring intensive care.

## Materials and Methods

### Study design and participants

Babies were recruited from the neonatal intensive care unit at Addenbrooke’s Hospital, Cambridge, UK. The study applied a randomized, open-label, single-center, two-arm, parallel design. Ethics approval was obtained prior to start of study recruitment. Eligibility criteria included birth weight less than 1200g and age less than 48h. Babies were excluded if they had a major congenital malformation or an underlying metabolic disorder, or mothers had had pregnancies complicated with diabetes. Informed written parental consent was obtained prior to study procedures.

All babies had real-time continuous glucose monitoring inserted within 48h post birth, which remained in situ for up to seven days. The paper-based guideline advised on the use of insulin or additional dextrose support. For a pre-specified period of 24h, between 48 and 72h post birth, a closed loop system controlled glucose in babies during the closed loop intervention, whereas babies in the control group continued to use real-time continuous glucose monitoring alongside the paper guideline to direct insulin therapy to maintain glucose control.

### Randomization

Babies were randomized within 48h of birth to closed loop or real time continuous glucose monitoring with a paper based guideline for insulin therapy. Randomization applied the minimization methods using the Minim randomization software.(*23*) Randomization was stratified according to gestational age and birth weight to ensure balance between the two groups.

### Common study procedures

Apart from glucose control over the pre-specified period between 48 and 72 hours, all other aspects of care including nutritional management were identical between treatment groups. Blood glucose monitoring was taken on the point of care glucose meter Statstrip® meter (Nova Biomedical, Waltham, MA, USA). Actrapid insulin (Novo Nordisk, Bagsværd, Denmark) in concentration of 25U/kg in 50 ml of 0.9% saline was used in both treatment groups. Study related activities were carried out until the end of the first week of life.

### Continuous glucose monitoring

An Enlite^TM^ sensor (Medtronic, Watford, UK) was inserted by hand into the lateral thigh of each baby and linked to the Paradigm® Veo^TM^ (Medtronic, Watford, UK) for calibration and to display the sensor glucose concentration. Real-time continuous glucose monitoring data were used by the clinical team caring for each baby. Nurses calibrated the sensor at least once every 12 hours with a blood glucose measurement taken on the Statstrip® meter (Nova Biomedical, Waltham, MA, USA) point of care glucose meter.

### Paper algorithm

The paper algorithm for insulin delivery was developed for the purposes of studies using real-time continuous glucose monitoring to optimize glucose control in preterm babies. It provided guidance based on the absolute sensor glucose value as well as glucose trends. If sensor glucose levels were outside of the target range or demonstrated a persistent trend the advice given was to review the clinical care and consider the need to check blood glucose level, review lines and nutritional intake and to consider modifying insulin delivery dose or providing additional dextrose. The bedside nurse could initiate a physician prescribed alteration in insulin delivery based on the paper algorithm. The insulin and dextrose were delivered by Alaris pumps (Carefusion, San Diego, CA, USA).

### Closed loop glucose control between 48 and 72 hours

Babies randomized to closed loop therapy were treated between 48 to 72 hours post birth using a closed loop system comprising (i) Enlite^TM^ sensor, (ii) a laptop computer running a model predictive control algorithm, and (iii) two Alaris syringe pumps. We used a control algorithm based on the model predictive control approach(*15*), optimized and tuned *in silico* using a computer simulation environment validated for glucose control in the critically ill,(*22*) and aiming to keep sensor glucose between 4.0 and 8.0 mmol/l. The algorithm calculated insulin or, at low glucose values, 20% dextrose infusion requirements based on real-time sensor glucose values. A study nurse entered sensor glucose values into the laptop and modified insulin and dextrose pumps as directed by the control algorithm every 15 minutes. During the closed loop intervention actrapid insulin in concentration of 5U/kg in 50 ml of 0.9% saline was used. The algorithm calculations utilized a compartment model of glucose kinetics(*24*) describing the effect of insulin on sensor glucose excursions. The algorithm was initialized using a baby’s weight and adapted itself to a particular baby by updating two model parameters – a rapidly changing glucose flux correcting for errors in model-based predictions and a slowly changing estimate of an insulin rate to maintain normoglycaemia. The individualized model forecasted plasma glucose excursions over a 1.5h prediction horizon when calculating the insulin rate and a 30 to 40 minute horizon when calculating the dextrose rate. Information about enteral or parenteral nutrition was not provided to the algorithm. A reference blood glucose value was used every 6h for calibration of glucose sensor. If sensor readings were not available due to sensor failure or loss of data capture, then hourly blood glucose levels were used to inform the algorithm for up to 4h. At this time, the algorithm continued to provide advice every 15minutes.

### Assessments and data collection

Demographic and clinical characteristics were collected at study initiation. Antenatal variables were defined as: antenatal steroids as having received at least one dose prior to delivery, prolonged rupture of membranes as rupture >24 hours prior to delivery, maternal smoking included mothers who smoked at any time during pregnancy and hypertension and chorioamnionitis were based on diagnoses recorded in the maternal medical file. All blood glucose measurements, insulin administration, type and volume of enteral and parenteral nutrition and additional intravenous glucose administration were recorded from the time of randomization to the end of continuous glucose monitoring.

### Statistical analysis

Investigators agreed on the outcome measures and the statistical analysis plan in advance. The primary outcome was the time spent in target glucose range between 4.0 and 8.0mmol/l as recorded by sensor glucose measurements. This data was compared between study arms. Secondary outcomes were time spent with sensor glucose levels between 2.6 and 10.0 mmol/, prevalence of hyperglycaemia (percent time sensor glucose >10.0 mmol/l), mean and standard deviation of sensor glucose. Safety endpoints included frequency of significant hypoglycaemia (any blood glucose <2.6 mmol/l) and other adverse events. As this was a feasibility study, no formal power calculations were performed. All analyses were performed on intention to treat basis. Unpaired t-test was used to compare normally distributed variables. Non-normally distributed variables were compared using Mann-Whitney U-test. Calculations were carried out using SPSS Version 23 (IBM Software, Hampshire, UK). Values are given as mean, standard deviation (SD) or median (interquartile range). P value less than 0.05 was considered statistically significant.

## Acknowledgments

The authors wish to acknowledge and thank all the families that took part in the study the clinical team on the Neonatal Unit and Prof D Rowitch for their support. We would also like to acknowledge the support of the Evelyn Trust, the National Institute of Health Research EME Program and the National Institute of Health Research Cambridge Biomedical Research Centre without whom the studies would not have been possible. We further thank Medtronic for providing the continuous glucose monitoring system and sensors for use in the study.

## Funding

Funding was provided by the Evelyn Trust, the National Institute of Health Research EME Program and the National Institute of Health Research Cambridge Biomedical Research Centre.

Medtronic provided the continuous glucose monitoring system and sensors. Medtronic had no role in design of the study, the gathering of data, access to data, preparation of the manuscript or decision to publish the results.

## Author contributions

LT contributed to the design of the study, to the acquisition of the data and drafting of the manuscript. DE contributed to the analysis of the data collected during the study, and critically reviewed the manuscript. RH contributed to the design of the study and in particular the model predictive computer algorithm, to the interpretation of the data, critical revision of the initial drafts of the manuscript. DBD contributed to the design of the study, evaluation of the data, critical review and revision of the manuscript. KB was the PI who had the original idea for the study and contributed to the study design, to the acquisition and analysis of the data prior to drafting the initial manuscript and subsequent critical revisions. All authors have approved the final manuscript prior to submission and are accountable for the integrity of the study.

